# HiC-SCA: A spectral clustering method for reliable A/B compartment assignment from Hi-C data

**DOI:** 10.1101/2025.09.22.677711

**Authors:** Justin Chan, Hidetoshi Kono

## Abstract

A/B compartment analysis identifies regions of active and inactive chromatin organization from Hi-C data. We present HiC Spectral Compartment Assignment (HiC-SCA), a graph-based method that models chromatin as a weighted network and uses spectral clustering to partition chromosomes into A and B compartments.

HiC-SCA includes four key improvements: a noise filter that removes low-quality data to prevent erroneous results, an orientation assignment method that correctly identifies A versus B compartments, a metric for selecting and assessing confidence in compartment assignments, and a resolution selection approach that identifies the highest resolution at which analysis can be performed for a dataset.

Evaluation across 21 Hi-C datasets from diverse cell types using cross-dataset correlation demonstrates that HiC-SCA achieves superior consistency compared to an established method, producing more reproducible compartment assignments between datasets of the same cell type.

HiC-SCA addresses key challenges in current A/B compartment analysis and provides researchers with robust tools for analyzing Hi-C datasets with varying quality.

## Introduction

Chromatin is the complex of DNA and proteins that packages genomic DNA within the nucleus of a cell. Understanding chromatin organization is fundamental to deciphering gene regulation mechanisms. While direct and indirect regulatory factors such as transcription factors and signaling pathways are well-characterized, the molecular mechanisms driving gene expression and repression through chromatin structure remain poorly understood. To address this knowledge gap, chromosome conformation capture techniques, particularly Hi-C^1–3^, have revolutionized our ability to study chromatin organization. Hi-C measures contact preferences or frequencies between all pairs of chromosomal loci, generating intra-chromosomal contact maps.

These contact maps reveal that chromatin structure is highly dynamic. Despite this inherent dynamism, intra-chromosomal contact maps consistently exhibit characteristic plaid-like patterns that indicate two primary groups of genomic loci (A- and B-compartments) that preferentially interact within their respective groups rather than between groups. A-compartments are associated with actively transcribed genes, while B-compartments correspond to repressed genes and heterochromatic regions^1^.

The most prevalent analytical approaches for identifying these compartments employ principal component analysis (PCA). These methods either compute correlations between contact patterns of all pairs of chromosomal loci to generate a correlation matrix subjected to eigen-decomposition or directly apply eigen-decomposition to the Hi-C contact matrix itself. The signs of the first or second eigenvector values determine compartment assignments. Examples of these approaches include POSSUMM^4^, FAN-C^5^, HiCExplorer^6^, Cooltools^7^, HOMER^8^, and dcHic^9^.

Alternatively, graph-based approaches, such as 4D Nucleosome^10^, model chromatin interaction data as a weighted graph and formulate compartment identification as a graph partitioning problem. Other graph-based methods like SCI^11^ use graph embedding. For more thorough reviews of these methods see Kalluchi et al.^12^ and Raffo and Paulsen^13^.

Current A/B compartment analysis methods face several limitations that affect assignment quality and consistency. Hi-C contact maps suffer from sparsity due to limited sequencing coverage, necessitating binning of loci into chromosomal intervals of fixed size, thereby determining the analysis resolution. However, current approaches rely primarily on sequencing coverage to estimate the appropriate resolution without considering data quality.

Additionally, coverage and data quality are not uniform across the same chromosome. This non-uniform coverage creates challenges, as low-coverage regions with few contacts may be incorrectly identified as a separate compartment. While matrix rebalancing methods^2,14^ address this by assuming uniform coverage across chromosomal intervals, this approach modifies the distribution of contacts. Alternative approaches that define individual genes as bins^10^ reduce sparsity but could exclude important regulatory regions.

Furthermore, eigen-decomposition based-methods suffer from orientation ambiguity in distinguishing A versus B compartments. This is typically resolved through incorporation of external knowledge such as gene density or GC content^12^. However, genes can transition between compartments depending on cell type and experimental conditions.

Perhaps most importantly, existing methods lack criteria for evaluating compartment assignment quality from Hi-C data itself, instead requiring validation through correlation with external datasets such as RNAseq gene expression or histone modification ChIP-seq data.

Here we present HiC Spectral Compartment Assignment (HiC-SCA, **Fig. 1**), a graph-based method that models chromatin (**Fig. 1a**) as a weighted network and uses spectral clustering to generate eigenvectors that partition chromosomes into A and B compartments. HiC-SCA addresses the identified problems through several key components. To improve data quality, HiC-SCA employs a low-coverage filter that removes noisy chromosomal intervals. We also introduce the inter-AB score, a novel metric that both identifies the optimal eigenvector for compartment partitioning and quantifies assignment quality from the eigenvector and eigenvalue. Using the selected eigenvector, the orientation selection algorithm determines the correct A and B compartment assignments using only Hi-C intra-interval contacts. HiC-SCA also incorporates a resolution selection method by assessing compartment assignment consistency across resolution scales. Through systematic performance evaluation using Matthews Correlation Coefficient analysis to compare across datasets of various cell types, we demonstrate that HiC-SCA achieves better assignment consistency compared to the established POSSUMM method.

**Fig. 1:**
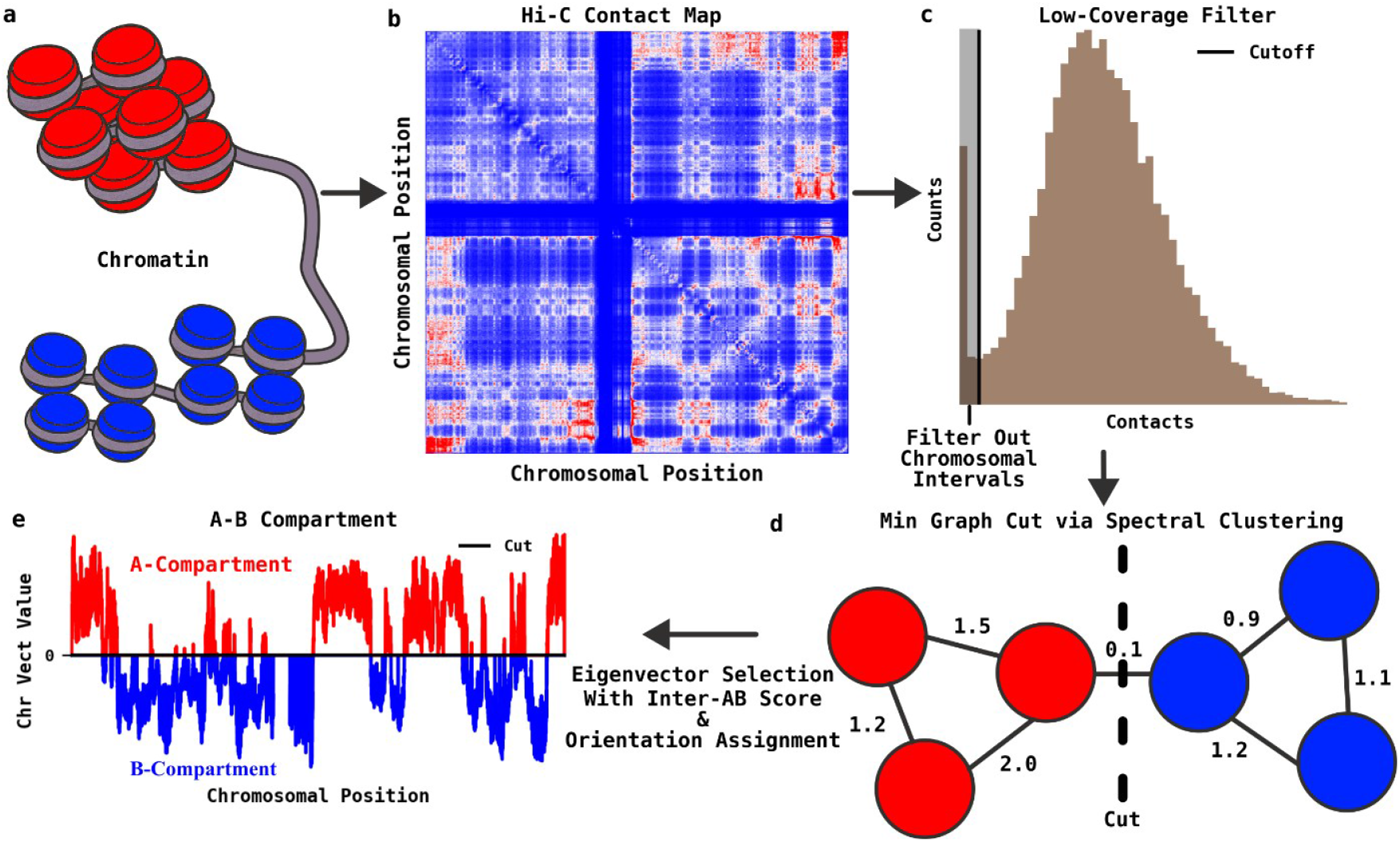
Overview of HiC Spectral Compartment Assignment (HiC-SCA). **a**, Chromatin organizations for A (red) and B (blue) compartments. **b**, A typical Hi-C map. Hi-C contact maps capture the organization of chromatin in the nuclei of cells. Raw Hi-C reads are binned into chromosomal intervals at a specific resolution (bin size) and normalized by expected background counts based on chromosomal interval distance to generate observed/expected (OE) contact maps. **c**, The distribution of average OE contacts for each chromosomal interval. Low-coverage chromosomal intervals are filtered from the OE contact map to reduce noise. **d**, A schematic view of a graph representing chromatin organization. Chromatin is modeled as a graph where each chromosomal interval represents a vertex, with edges weighted by the filtered OE contact values. The graph is partitioned into two compartments by finding cuts that minimize edge weights. Spectral clustering provides approximate solutions to this minimal cut problem, with each eigenvector representing a different partitioning solution. The optimal eigenvector for A/B compartment assignment is selected using the inter-AB score, and its orientation is determined by analyzing intra-interval contact patterns. **e**, A/B compartment assignment from the chromosomal vector. The chromosomal vector, where the eigenvector values are mapped to their respective chromosomal interval position. Chromosomal intervals are assigned to A-compartments if eigenvector values >0, B-compartments if the values <0, or unassigned if values = 0.

## Results

### Overview of HiC-SCA

Hi-C data analysis begins with binning raw sequencing reads into chromosomal intervals at a specified resolution (e.g. 1kb, 5kb), where each interval represents a contiguous genomic region of fixed size. Contact counts between intervals are normalized by expected background counts based on genomic distance to generate observed/expected (OE) contact maps (**Fig. 1b**), thereby correcting for the general decay of contact frequency with increasing chromosomal distance. HiC-SCA models each chromosome as a weighted graph where chromosomal intervals serve as vertices (nodes) and edges (links) are weighted by the OE contact values (**Fig. 1d**).

The A/B compartment assignment problem is formulated as finding the minimum graph cut that partitions the chromosome into two compartments with minimal inter-compartment contacts. This optimization problem can be reformulated into the graph Laplacian form, which closely resembles the spectral clustering problem. Spectral clustering provides an approximate solution to the minimum cut problem through eigen-decomposition of the Laplacian matrix. Each eigenvector represents a different partitioning solution, with positive and negative values corresponding to the two compartments respectively. However, this approach faces three related challenges: solutions with the smallest cuts may represent artifacts arising from low-coverage chromosomal intervals, the approximation means the eigenvector corresponding to the smallest cut may not represent the optimal solution and the sign of eigenvector values is arbitrary, requiring determination of which compartment represents A versus B.

To address these issues, HiC-SCA employs a low-coverage filter (**Fig. 1c**) that removes noisy intervals, thereby reducing the likelihood of artifact solutions. Additionally, we introduce the inter-AB score that evaluates assignment confidence, enabling selection of the optimal eigenvector for A/B assignment. Finally, the correct orientation of the selected eigenvector is determined to distinguish A from B compartments (**Fig. 1d**) using intra-interval contact patterns.

### Performance Evaluation of HiC-SCA

A/B compartment assignment methods are commonly validated by correlating assignments with gene expression data and histone post-translational modifications (PTMs) that serve as markers for transcriptional activity or repression, based on the assumption that A-compartments correspond to transcriptionally active euchromatic regions while B-compartments represent repressed heterochromatic regions^1^. However, this approach has significant practical constraints: it requires gene expression and chromatin modification data obtained under identical experimental conditions as the Hi-C data, and since these relationships are correlative, they may not hold consistently across all experimental conditions. These limitations motivated the development of an alternative evaluation framework that assesses compartment assignment quality by evaluating whether assignments are reproducible across datasets.

We evaluate the performance of HiC-SCA by assuming that assignments should be more similar between datasets from the same cell type than between different cell types. Matthews Correlation Coefficient (MCC) was employed to quantify agreement between compartment assignments from different datasets for each chromosome, as MCC tolerates uneven compartment sizes and outliers.

Analysis of 21 Hi-C datasets representing diverse human cell lines and types demonstrates that HiC-SCA achieves the expected performance pattern (**Fig. 2a**). Same cell types exhibit higher correlation values than comparisons between different cell types, with notable exceptions for myelogenous leukemia cells, datasets generated with the older solution Hi-C protocol or those with low sequencing coverage. Comparison with POSSUMM, a commonly used PCA-based method, reveals that POSSUMM shows similar overall patterns but achieves lower correlation values (**Fig. 2b**). Direct comparison between the two methods confirms that datasets where POSSUMM demonstrates high cross-dataset correlation also show strong agreement with HiC-SCA assignments (**Fig. 2c**). Since POSSUMM represents an independent method previously validated against gene expression and chromatin modification data, this concordance indicates that HiC-SCA’s consistent assignments reflect genuine compartmentalization. This suggests our method is detecting real biological signals rather than artifacts.

**Fig. 2:**
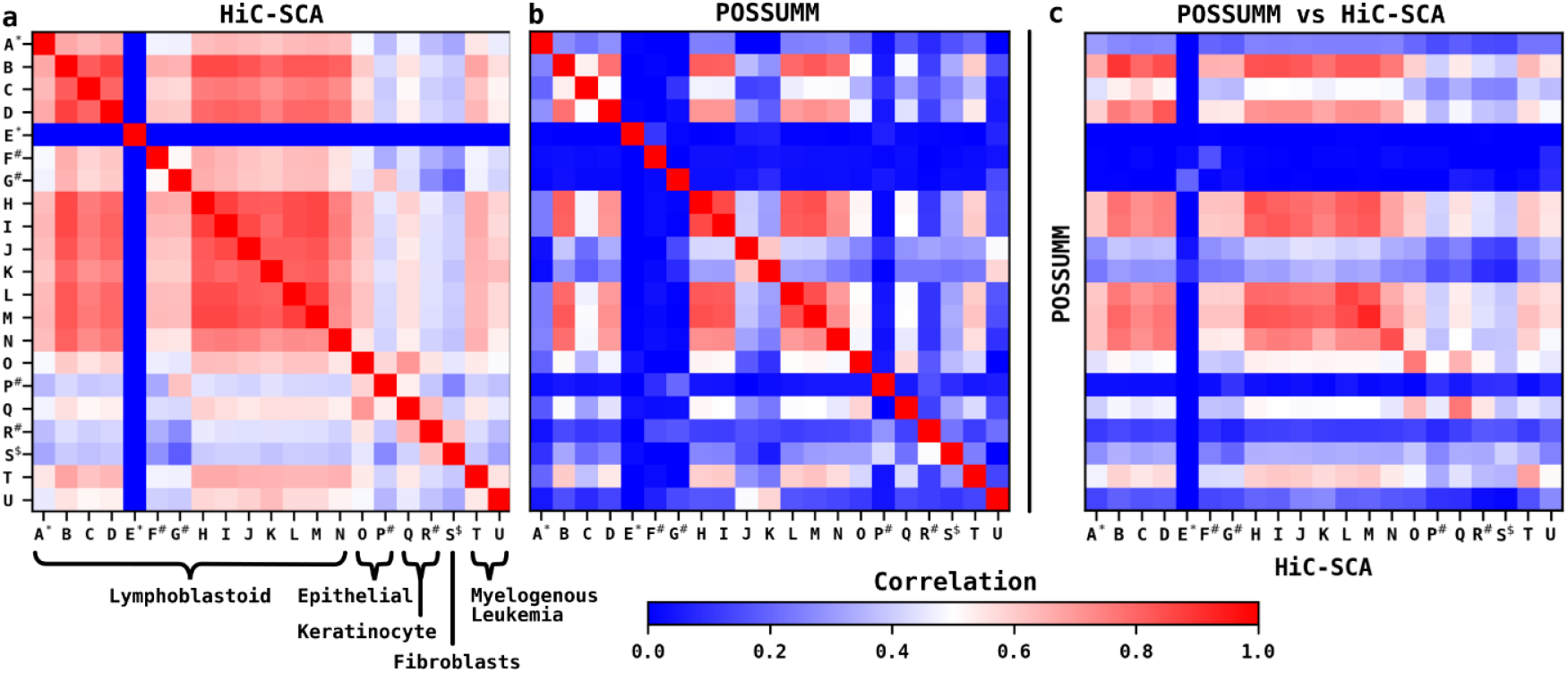
Performance of HiC-SCA vs POSSUMM. **a**, Cross-dataset Matthews Correlation Coefficients (MCC) matrix for HiC-SCA. HiC-SCA performance was evaluated using cross-dataset correlation analysis based on the principle that A/B compartment assignments should be more consistent between datasets from the same cell type than between different cell types. MCC were computed between all pairs of 21 Hi-C datasets (A-U, **Supplementary Table 1**) representing various human cell lines and cell types. Datasets marked with “#” were obtained using solution Hi-C protocol (older and lower quality protocol) while unmarked datasets were obtained using in-situ Hi-C protocol (newer and higher quality protocol). Datasets with “*” have low coverage (< 100m intra-chromosomal reads), and datasets with “$” were obtained using solution Hi-C protocol with low coverage. Results show the expected pattern of higher correlations within cell types, with exceptions for low-coverage datasets and some solution Hi-C datasets. **b**, Cross-dataset MCC matrix for POSSUMM. Identical analysis performed using POSSUMM, a commonly used method for A/B-compartment assignment that performs PCA on the correlation matrix of contact patterns between chromosomal intervals. The first principal component from POSSUMM’s analysis is used for compartment assignments. POSSUMM shows lower consistency in assigning similar A/B compartments between datasets from the same cell type. **c**, Cross-correlation matrix of A/B compartment assignments between HiC-SCA and POSSUMM across all dataset pairs. Datasets where POSSUMM assignments are consistent across datasets of the same cell type also show high agreement with HiC-SCA. This demonstrates that HiC-SCA not only provides consistent compartment assignments but also produces biologically meaningful results that align with an independent method. **a-c**, Limited to the 22 autosomal chromosomes, performed at 5 kb resolution. The correlation values represent absolute correlation while the sign corresponds to orientation similarity or difference. See **Supplementary Fig. 1** for cross-dataset MCC at other resolutions.

### Low Coverage Filter

The minimum graph cut formulation for A/B compartment assignment can produce trivial solutions^15^ when chromosomal intervals exhibit low coverage and sparse interactions with the rest of the chromosome (**Fig. 3a**). In these cases, poorly connected intervals are isolated into a single small compartment while all remaining intervals are grouped into one large compartment, resulting in biologically meaningless partitions (**Fig. 3b**).

**Fig. 3:**
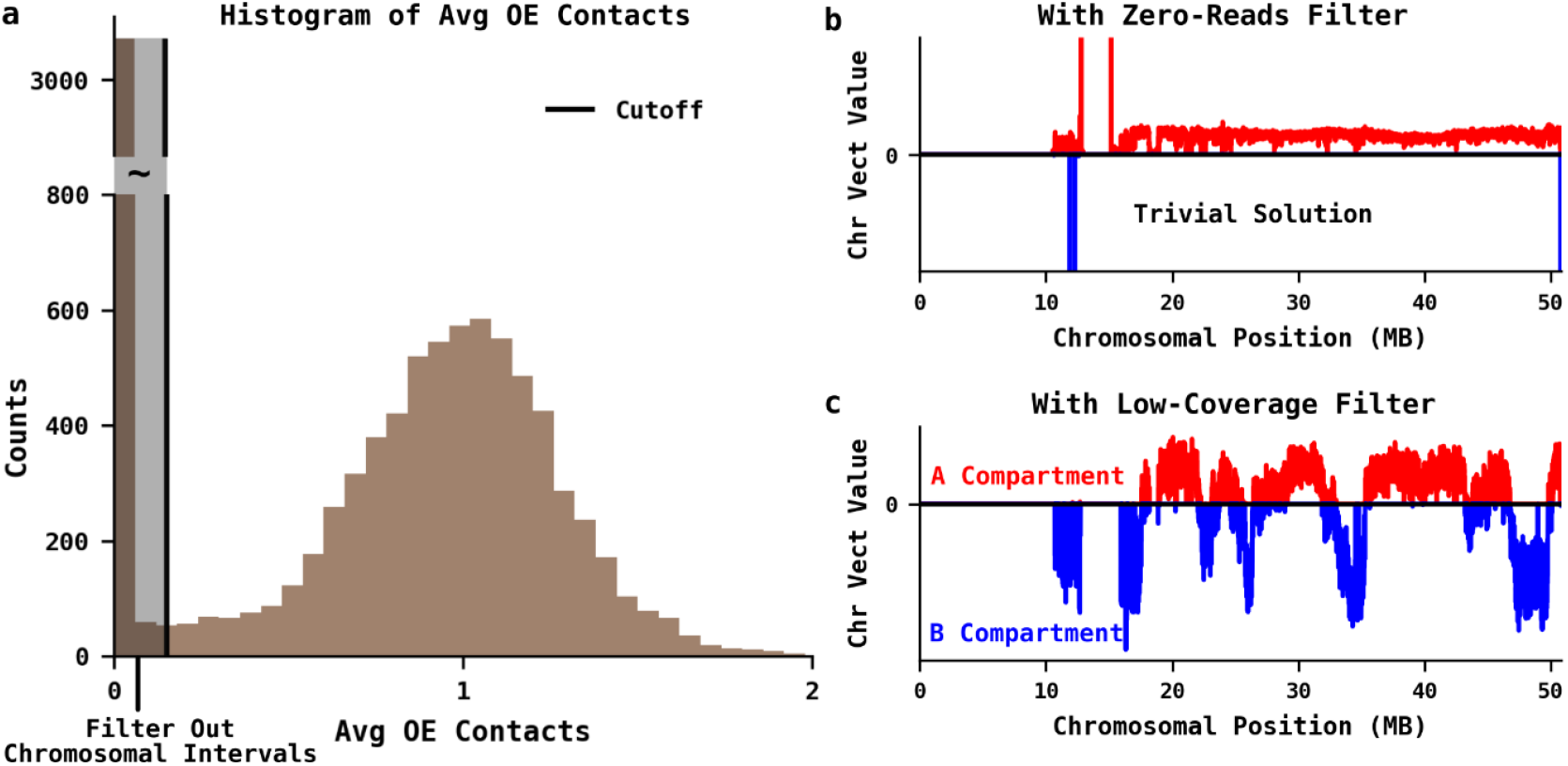
Low-coverage filter. **a**, Distribution of average observed-over-expected (OE) contact values and filter threshold identification. The distribution of average OE contact values for each chromosomal interval typically shows a bimodal pattern. The low-coverage filter identifies both peaks and uses the minimum between them as a cutoff threshold. Chromosomal intervals below this cutoff are excluded from downstream analysis, with intervals having non-zero contact values automatically assigned to B-compartment. **b**, Chromosomal vector solution after removing only intervals with zero contact. Without applying the low-coverage filter, spectral clustering suffers from a common problem where poorly connected chromosomal intervals are isolated into their own small compartment while the remaining intervals form a single large compartment, producing trivial solutions. **c**, Chromosomal vector solution after low-coverage filtering. Applying the low-coverage filter prior to spectral clustering reduces the occurrence of these artifacts and enables the identification of biologically relevant A/B compartments.

To address this issue, HiC-SCA employs a low-coverage filter that identifies and removes problematic intervals prior to spectral clustering. The filter analyzes the distribution of average OE contact values for each chromosomal interval, which commonly exhibits a bimodal pattern with two distinct peaks (**Fig. 3a**). The algorithm identifies both peaks and determines the minimum between them as a cutoff threshold. Chromosomal intervals with average contact values below this threshold are excluded from the contact map used for spectral clustering, while any excluded intervals with non-zero contact values are automatically assigned to the B-compartment. Application of this filtering approach prior to spectral clustering reduces the occurrence of trivial partitioning artifacts and enables the identification of biologically relevant A/B compartments (**Fig. 3c**).

### Orientation Assignment using Intra-chromosomal Intervals

The orientation of eigenvectors obtained from PCA-based methods and spectral clustering is inherently arbitrary, as flipping the sign of an entire eigenvector (multiplying by −1) yields an equally valid mathematical solution, thus presenting a challenge for distinguishing A from B compartments. Currently, POSSUMM addresses this by comparing solutions against pre-assigned 100 kb resolution references for human cells, selecting the orientation with better agreement. However, this approach relies on prior biological knowledge and may not generalize across different cell types or experimental conditions.

HiC-SCA determines compartment orientation directly from intrinsic properties of the Hi-C data rather than external references. The intra-interval contacts (diagonal elements of the OE contact map, **Fig. 4a**) exhibit a bimodal Gaussian-like distribution (**Fig. 4b**). Each Gaussian component corresponds to one of the two compartments, with the higher mean consistently representing A-compartments and the lower mean representing B-compartments (**Fig. 4c**). While these Gaussian components show substantial overlap that limits their utility for assigning individual intervals to compartments, they provide a reliable signal for orientation determination by identifying which compartment exhibits higher average intra-interval contacts.

**Fig. 4:**
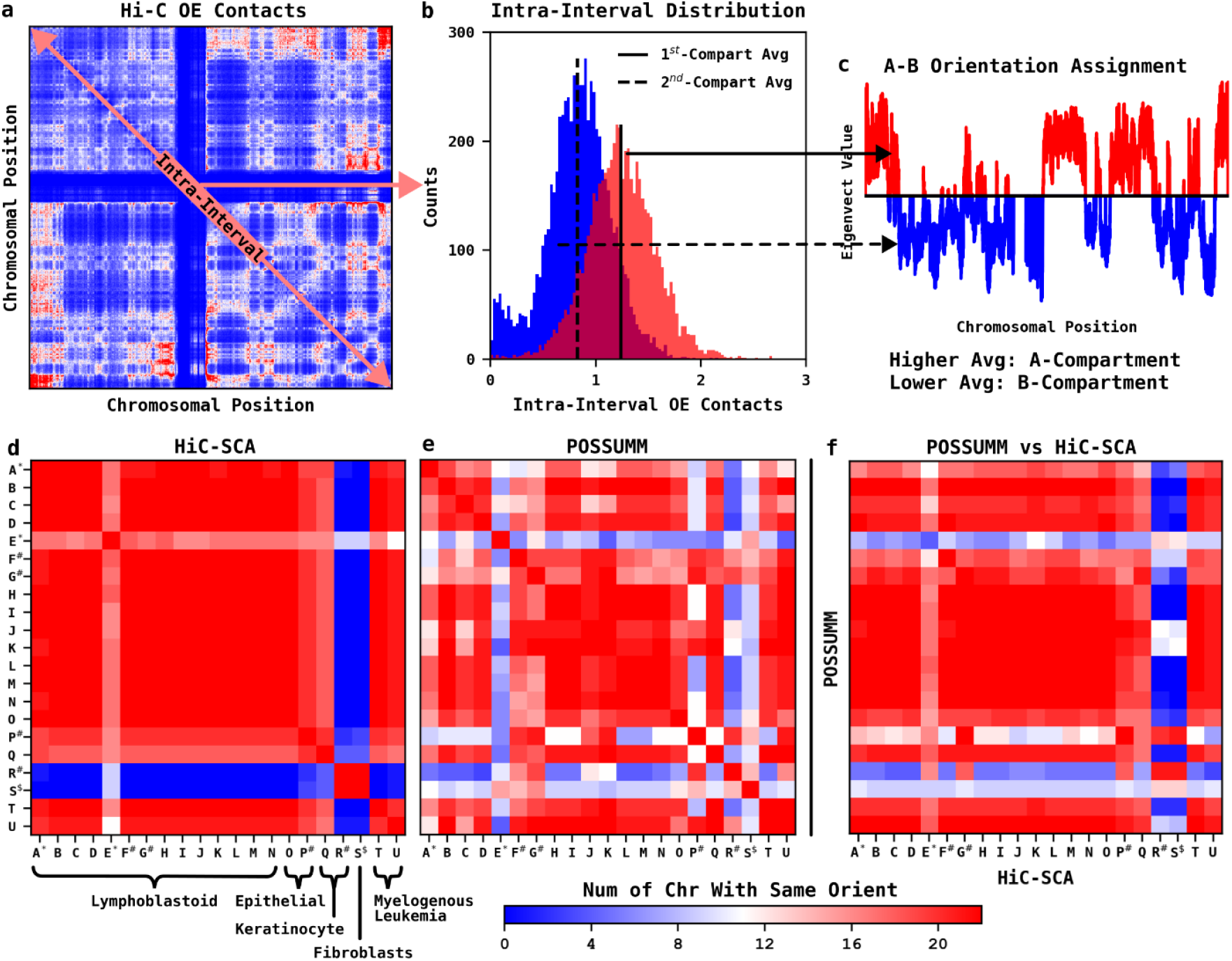
Orientation Assignment. **a**, Hi-C OE contact map with intra-interval contacts. The diagonal elements represent intra-interval OE contacts used for orientation determination. **b**, Distribution of intra-interval OE contact values. The intra-interval OE contacts show a bimodal Gaussian-like pattern. A-compartments tend to be represented by the Gaussian with the higher mean and B-compartments by the Gaussian with the lower mean. **c**, Orientation selection based on intra-interval contact distribution. This bimodal pattern is exploited to determine the correct orientation of the arbitrarily signed eigenvector from spectral clustering. **d**, Cross-dataset orientation agreement matrix for HiC-SCA (chromosome counts). Orientation agreement between HiC-SCA assignments across dataset pairs, measured as the number of chromosomes with consistent orientation (positive MCC). Results show high orientation agreement except for low-coverage dataset E and solution Hi-C datasets R and S. **e**, Cross-dataset orientation agreement matrix for POSSUMM (chromosome counts). Identical analysis for POSSUMM shows overall high orientation consistency but lower than HiC-SCA. **f**, Orientation agreement matrix (chromosome counts) between POSSUMM and HiC-SCA. Direct comparison demonstrates high overall concordance in A/B compartment orientation assignments, confirming that the distribution patterns in **b** are biologically meaningful. **d-f**, Dataset labeling follows **Fig. 3** conventions. Analysis performed on 22 autosomal chromosomes at 5 kb resolution. See **Supplementary Fig. 2** for orientation agreement (chromosome counts) at other resolutions.

We evaluated orientation assignment consistency by counting chromosomes with identical MCC signs between dataset pairs across all 21 Hi-C datasets. HiC-SCA demonstrates exceptionally high orientation agreement across datasets regardless of cell type, with notable exceptions only for solution Hi-C datasets R and S (**Fig. 4d**). Even the low-coverage dataset E maintains substantial agreement with most other datasets, indicating robust orientation assignment even under suboptimal conditions. Comparison with POSSUMM reveals that while POSSUMM achieves generally high orientation agreement, it exhibits lower consistency than HiC-SCA (**Fig. 4e**). Direct comparison between the two methods confirms high concordance in orientation assignments, validating the observed distribution patterns underlying HiC-SCA’s approach (**Fig. 4f**). These results demonstrate that HiC-SCA’s orientation assignment method achieves high consistency while relying exclusively on intrinsic Hi-C data features.

### Inter-AB Score for Eigenvector Selection and A/B-Compartment Assignment Confidence Assessment

The spectral clustering formulation underlying HiC-SCA provides additional insights beyond the original minimum graph cut problem. These insights can be used for both eigenvector selection and assignment confidence assessment. In the graph Laplacian formulation, each eigenvalue represents the sum of squared differences between interval values in the corresponding eigenvector, weighted by their OE contact values. These weighted differences can be decomposed into intra-compartment and inter-compartment components. While the original discrete minimum graph cut problem seeks solutions with constant values for each compartment with zero intra-compartment differences, the relaxed spectral clustering formulation allows for variable intra-compartment and inter-compartment differences that reflect the underlying graph topology.

The eigenvector element values provides information about interval connectivity that extends beyond simple compartment assignment. Intervals with values close to zero are more connected to both compartments, indicating ambiguous assignment, while intervals with values that are further away have strong preferential connections to one compartment with few or weak connections to the other.

We exploit these relationships through the inter-AB score, which quantifies the ratio of the average inter-compartment difference to the average background difference. The inter-AB score is calculated as the weighted average of squared inter-compartment differences divided by the normalized eigenvalue, where the normalized eigenvalue represents the weighted average of all squared differences between chromosomal interval pairs and serves as a measure of background difference levels.

This formulation provides a signal-to-noise ratio that measures compartment separation confidence (**Fig. 5a**).

**Fig. 5:**
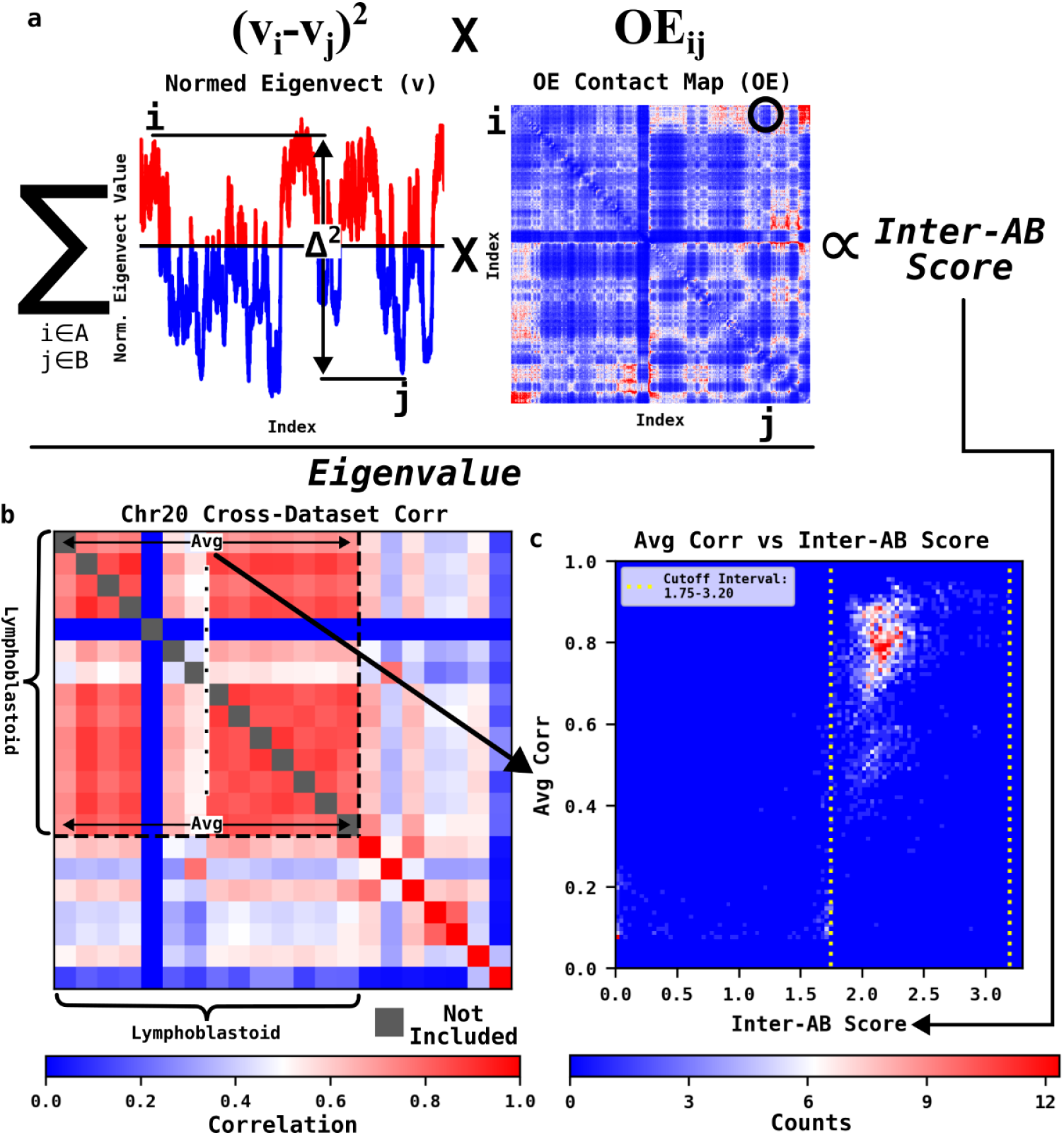
Inter-AB score for assessing compartment assignment confidence. **a**, Definition and computation of inter-AB score. The inter-AB score quantifies how well spectral clustering separates the two compartments, with higher scores indicating greater confidence in assignment confidence. The score is proportional to the sum of squared differences between the normalized eigenvector values of intervals from different compartments (positive/negative values), weighted by their respective OE contact values and normalized by the eigenvalue. See Methods for exact computation details. **b**, Example chromosomal cross-dataset MCC matrix and average correlation computation. To establish confidence thresholds for inter-AB scores, we calibrate them against average correlation between lymphoblastoid cell datasets (A-N), which should have highly similar A/B compartments. Cross-dataset MCC matrix for Chr20 at 5 kb resolution is shown as an example demonstrating how average correlation is computed. The average correlation serves as a confidence score, as differences between these similar datasets can be attributed to assignment confidence. **c**, Heatmap of inter-AB scores versus average correlations. Each count represents a specific chromosome, resolution, and dataset combination. Inter-AB scores between 1.75 and 3.2 reliably identify high-confidence A/B compartment assignments (average correlation > 0.5).

The inter-AB score serves dual purposes in HiC-SCA: eigenvector selection and assignment confidence assessment. For eigenvector selection, a modified version balances cut size with compartment separation confidence. Among the computed eigenvectors, the first (trivial) eigenvector is excluded as it assigns all chromosomal intervals to a single compartment, representing a zero cut with no meaningful partition. Small eigenvalues approximate solutions with small cuts, representing partitions that remove low total edge weights from the graph. To incorporate both cut size and separation confidence, the inter-AB score is divided by the ratio of the current eigenvalue to the second-lowest eigenvalue. As the current eigenvalue increases relative to the second eigenvalue, this ratio increases, which linearly decreases the inter-AB score, thereby balancing preference for both small cuts and high-confidence separation. Among the 10 smallest non-trivial eigenvectors, the eigenvector with the highest modified inter-AB score is selected for compartment assignment.

For assignment confidence assessment, the inter-AB score requires calibration to establish meaningful interpretation thresholds. We calibrated the score using lymphoblastoid cell line datasets under the assumption that datasets from the same cell type should exhibit similar compartment assignments. We attributed any differences between lymphoblastoid datasets to variations in data quality and coverage. For each specific combination of dataset, chromosome, and resolution, we computed the MCC between that dataset and each of the other lymphoblastoid datasets for the same chromosome and resolution. We then averaged these values to generate a quality score for the corresponding A/B compartment assignment (**Fig. 5b**).

We define high-confidence compartment assignments as those with average correlations >0.5. Analysis reveals that inter-AB scores between 1.75 and 3.2 reliably predict high-quality assignments (**Fig. 5c**), with the upper bound established because scores above 3.2 typically represent problematic assignments. This creates a binary classification where compartment assignments are categorized as either high-confidence or low-confidence. Using average correlation as the reference and inter-AB score cutoffs as the classifier, we achieve a MCC of 0.75, showing that inter-AB scores can effectively identify high-confidence assignments.

These established thresholds enable users to assess compartment assignment confidence directly from their Hi-C data, providing a measure that complement traditional external validation approaches.

### Resolution Selection

Currently, there is no rigorous method for selecting the highest resolution at which meaningful A/B compartment analysis can be performed. The maximum resolution that a dataset can support depends on both sequencing coverage and data quality, yet existing approaches typically rely on sequencing coverage alone.

This limitation motivated the development of a systematic approach for resolution optimization based on the principle that if data supports analysis at higher resolution, the assigned compartments should remain consistent with those identified at lower resolutions.

The solution eigenvector from spectral clustering excludes filtered low-coverage intervals; we create a chromosomal vector by mapping these eigenvector values back to their respective intervals across the entire chromosome. To test consistency across resolutions, contiguous intervals in the higher resolution chromosomal vector are combined to predict compartments at lower resolutions using weighted averages, where the weights correspond to the squared values from the vector. The MCC between predicted and directly assigned compartments at lower resolutions indicates whether the higher resolution assignments accurately capture the underlying compartment patterns (**Fig. 6a**). This approach identifies the optimal resolution by determining the finest scale at which assignments remain predictive of coarser-scale compartments.

**Fig. 6:**
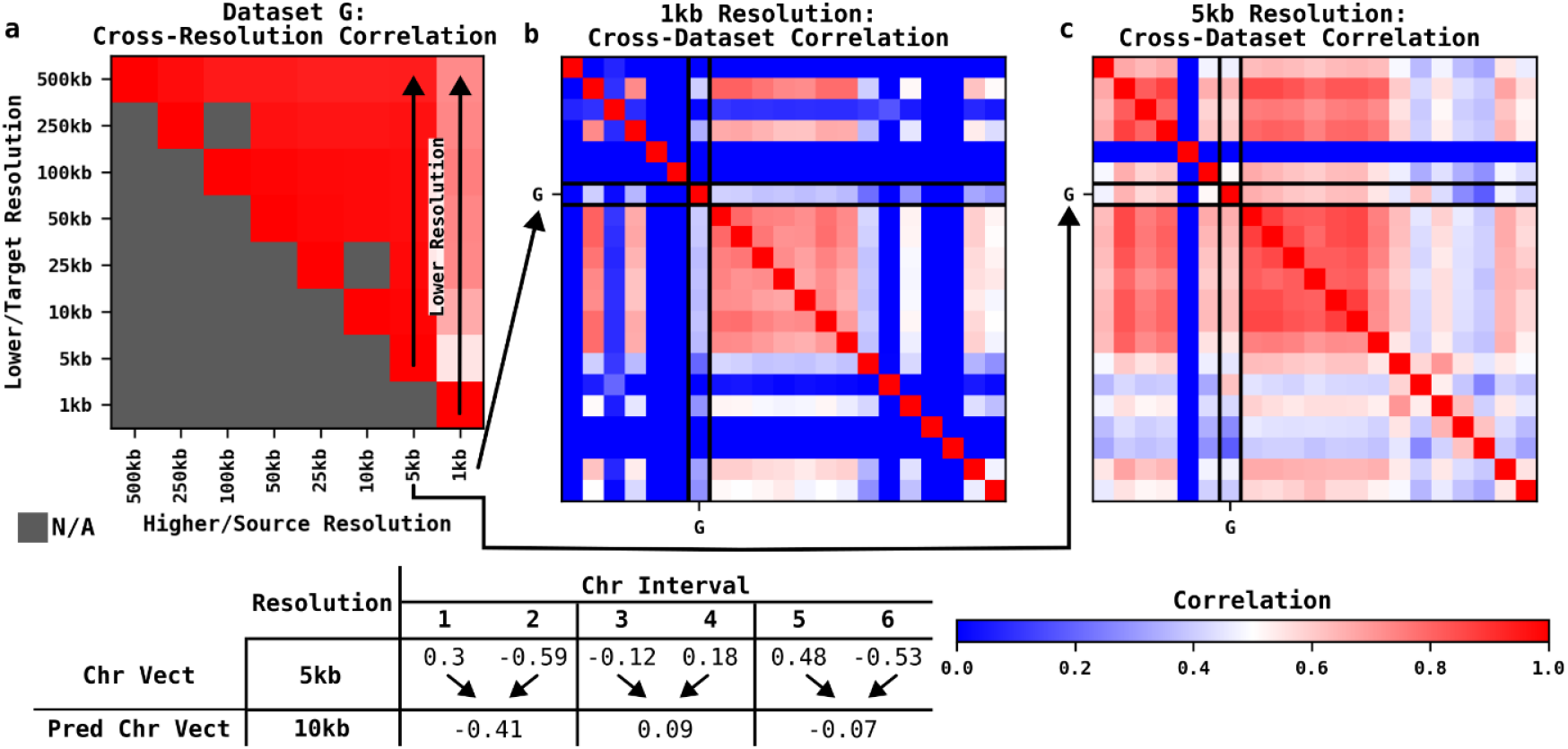
Resolution selection for A/B-compartment analysis. **a**, Cross-resolution MCC matrix for dataset G. To determine the optimal resolution for A/B-compartment analysis, assignments were performed at multiple resolutions and validated through cross-resolution prediction. Higher resolution assignments (smaller chromosomal intervals) were used to predict lower resolution compartments by computing weighted averages of chromosomal vector values, where the weights correspond to the squared chromosomal vector values that sum to one (see bottom table). The MCC between predicted and directly assigned compartments at lower resolutions indicates whether the higher resolution analysis can accurately reproduce coarser-scale compartments. Each column shows the current resolution’s prediction correlation with lower resolution assignments, while rows represent the ability of higher resolutions to reproduce the assignments at the current resolution. For dataset G, 1 kb resolution poorly reproduces lower resolution assignments (vertical arrow along 1 kb column), while 5 kb resolution (vertical arrow along 5 kb column) and lower resolutions show good reproduction. **b**, Cross-dataset MCC matrix at 1 kb resolution. Dataset G shows poor correlation with other datasets at 1 kb resolution. **c**, Cross-dataset MCC matrix at 5 kb resolution. At 5 kb resolution, dataset G shows improved agreement with other datasets. This analysis identifies the highest resolution that the dataset can support, but as seen with dataset G at 5 kb resolution (**Fig. 2a**, lower agreement with other lymphoblastoid datasets), it cannot evaluate assignment quality.

Applying this approach to dataset G demonstrates its effectiveness in resolution selection. Analysis at 1 kb resolution fails to reproduce compartment patterns at lower resolutions. However, 5 kb resolution and coarser scales show strong reproduction of lower resolution assignments, identifying 5 kb as the highest suitable resolution for this dataset. Cross-dataset correlation confirms this assessment, showing that dataset G exhibits poor agreement with other lymphoblastoid datasets at 1kb resolution (**Fig. 6b**) but substantially improved at 5 kb resolution (**Fig. 6c**). While this method successfully identifies the maximum supportable resolution, it cannot independently assess assignment quality, as evidenced by dataset G’s lower agreement with other lymphoblastoid datasets even at the optimal 5 kb resolution.

## Discussion

HiC-SCA addresses fundamental challenges in A/B compartment analysis through a methodological approach that reduces assumptions about Hi-C data characteristics while exploiting empirical patterns observed within the data itself. Rather than extensively modifying original Hi-C data through normalization or transformation, HiC-SCA preserves experimental data while identifying and excluding problematic regions based on their observable distribution patterns. This data-driven principle extends to compartment orientation determination, where HiC-SCA relies on empirical analysis of intra-chromosomal contact patterns rather than external prior knowledge.

### Matrix Rebalancing

The central challenge in Hi-C data analysis stems from two interrelated issues: the inherent sparsity of chromatin contact data and the non-uniform coverage across chromosomes. The sparsity necessitates binning reads into chromosomal intervals and careful determination of bin size. Additionally, some genomic regions are captured with higher coverage than others, creating biases in the contact matrix. Existing methods commonly employ matrix rebalancing strategies to address the non-uniform coverage problem by normalizing contact frequencies across all chromosomal intervals, operating under the assumption that all intervals should be equally visible in the contact map. However, HiC-SCA’s improved performance compared to POSSUMM, which employs matrix rebalancing, suggests that matrix rebalancing may be unnecessary for effective compartment analysis.

Furthermore, the justifications for matrix rebalancing remain problematic. While these methods claim to correct for technical biases introduced during sequencing or PCR amplification, the normalization procedures do not actually address the specific sources of bias they purport to correct. If the goal were genuinely to normalize for sequencing bias, the methods would need to account for factors such as GC content and specific sequence motifs. Instead, matrix rebalancing simply alters the contact distribution patterns to meet the constraint that each interval must have the same coverage, making strong assumptions about the relative contact frequencies that different chromatin regions should exhibit while obscuring natural distribution patterns in the data.

### Computational Efficiency

PCA-based approaches have attempted to circumvent the sparsity challenge in eigen-decomposition by computing cross-correlations between contact patterns of all chromosomal interval pairs, thereby generating dense correlation matrices. While this strategy addresses sparsity, it creates prohibitive memory requirements that limit analysis at higher resolutions. POSSUMM addresses this limitation by avoiding computing the full correlation matrix. In contrast, HiC-SCA’s low coverage filter and graph Laplacian formulation remains sparse and computationally efficient, enabling analysis of high-resolution datasets without excessive memory or computational demands.

### Scale of A/B Compartments

Our empirical analysis of cross-resolution consistency reveals important insights about the biological scale at which A/B compartments operate reliably. Our cross-resolution correlation analysis demonstrates that compartment assignments performed at higher resolutions begin to lose consistency with lower resolution assignments starting around 100 kb interval sizes (**Supplementary Fig. 3**). This pattern suggests that the biological scale at which A/B compartments function meaningfully is finer than 100 kb. Based on these findings, we recommend performing A/B compartment analysis at resolutions of 10 kb or finer to maintain sufficient resolution while providing a safety margin from the 100 kb threshold where consistency begins to deteriorate.

## Methods

### Obtaining and Parsing the Hi-C Datasets

Hi-C datasets (.hic files) were obtained from ENCODE portal^16^ when available. If the Hi-C data were not available, the raw high-throughput sequencing files (.fna files) were processed with Juicer 2 (pulled from repository with last commit on November 23, 2024)^17^. Raw sequencing datasets were aligned to the reference human genome assembly GRCh38.p14.

The 21 Hi-C datasets were from human cell lines reported by Rao et al.^2^ and Harris et al.^4^. See **Supplementary Table 1** for details.

Hi-C files were parsed and loaded into Python with hic-straw 1.3.1^18^ as sparse SciPy matrices.

### Computing the Observed/Expected (OE) Contact Map

The OE normalization removes distance-dependent contact decay patterns inherent in Hi-C data through a four-step procedure similar to the approach implemented in Juicer^17^. For each chromosome *c* with contact matrix M_*c*_ of size *n* × *n* chromosomal intervals (bins), we first calculated the background contact counts as a function of interval distance *d*:

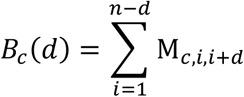

The number of interval pairs at chromosomal distance *d* is *N*_*c*_(*d*) *= n – d*.

To deal with the sparsity of counts, distances with fewer than 400 total contacts were smoothed out by extending the binning window from *d* to *d + k*, where *k* is the smallest integer such that the cumulative contact count reaches or exceeds 400. The smoothed background count is:

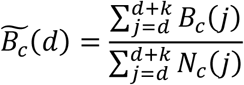

The genome-wide average background was computed by:

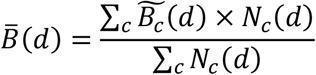

Since chromosomes exhibit different total contact counts, the genome-wide average background 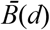 requires chromosome-specific correction factors. This factor is the ratio of observed to expected total contacts:

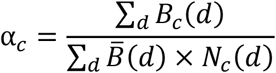

The final normalized contact matrix was obtained by dividing each entry in M_*c*_ by its chromosome-corrected expected background count:

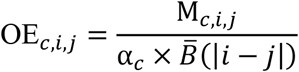

### Low Coverage Filter

Chromosomal intervals with low coverage and sparse interactions can produce trivial solutions during spectral clustering. We implemented a filtering procedure to identify and remove problematic intervals based on their average OE contact values.

For each chromosomal interval *i*, we calculated its average OE contact value as:

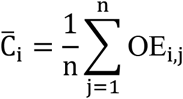

where *n* is the total number of intervals on the chromosome.

Since the distribution of average OE contact values typically exhibits a bimodal pattern, we constructed a histogram using 50 equally spaced bins from 0 to 3 and identified peaks by detecting gradient sign changes in the histogram. Peak locations were identified at positions where

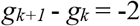

where *g*_*k*_ = sign(*h*_*k+1*_ - *h*_*k*_) and sign(x) returns +1, 0, or −1, and *h*_*k*_ represents the histogram count in bin *k*. Only peaks exceeding 1% of total intervals were retained.

For multiple peaks located at histogram bin positions *p*_*1*_, *p*_*2*_, …, *p*_*x*_, we identified the first peak *p*_*1*_ and the peak with the highest histogram count:

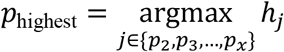

The cutoff was determined by finding the histogram bin with the minimum count between these two peaks:

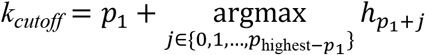

The midpoint average OE value of the *k*_*cutoff*_ bin (*b*_*cutoff*_) was selected as the threshold value, capped at a maximum of 0.5.

Chromosomal intervals *i* were retained if:

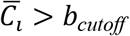

The procedure was applied iteratively up to three times until no additional intervals fell below the threshold. Filtered chromosomal intervals with non-zero average OE contacts are assigned to the B-compartment, while intervals with zero average OE contacts remain unassigned.

### Spectral Clustering for A/B Compartment Assignment

We applied spectral clustering to partition chromosomal intervals into A/B compartments by modeling each chromosome as a weighted graph where intervals serve as vertices and edges are weighted by OE contact values. The partitioning problem was formulated as a normalized cut (Ncut)^19^ optimization, a variant of the minimum cut problem that penalizes highly unbalanced partitions by normalizing the cut size by the number of intervals in each partition:

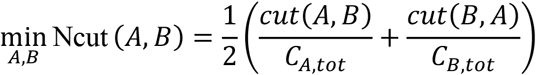

where cut(*A, B*) represents the total edge weights between compartments A and B. With *C*_*X*,tot_ = Σ_*i*∈*X*_ *C*_*i*_ denoting the total sum for all intervals in compartment *X* and *C*_*i*_ = Σ_*j*_ *OE*_*ij*_.

We solved this optimization problem using the symmetric normalized Laplacian L_sym_ = D^-1/2^LD^- 1/2^, where L = D - W is the unnormalized Laplacian and D is the diagonal degree matrix with D_*ii*_ = *C*_*i*_. The Ncut problem is equivalent to the quadratic form 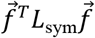, where 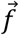 represents the solution vector with discrete assignment values. The original discrete optimization problem is NP-hard, requiring exhaustive search across all possible partitions. This problem can be approximated by relaxing the discrete assignment constraints to allow 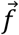 to take continuous values, transforming it into a spectral clustering problem solved through eigenvalue decomposition of L_sym_ (see von Luxburg^15^ for details).

We computed the 11 eigenvectors corresponding to the smallest eigenvalues of L_sym_, where 2^nd^- 11^th^ smallest eigenvectors serve as candidate solutions for compartment assignment, excluding the first trivial eigenvector. For each candidate eigenvector, intervals with positive values are assigned to one compartment and intervals with negative values to the other compartment, though the designation of which compartment represents A versus B is determined through orientation selection (see below).

### The Inter-AB Score

The *x*^th^ eigenvector 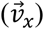 of L_sym_ is related to its eigenvalue (λ_x_) through the quadratic form by:

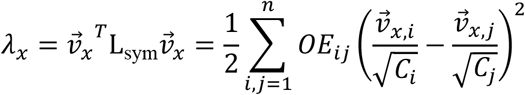

where 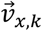, is the *k*^th^ element value in the eigenvector and 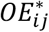 represents the filtered OE contact map.

This can be decomposed into:

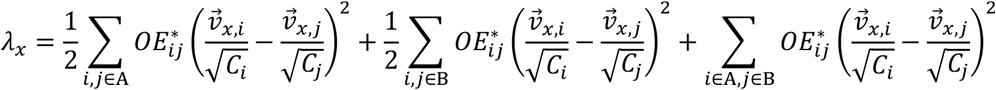

With the first two terms representing the intra-compartment weighted sum of squares difference while the last term represents the inter-AB weighted sum of squares difference.

Based on the decomposition above, we define the inter-AB score by first computing the weighted average of the inter-compartment term:

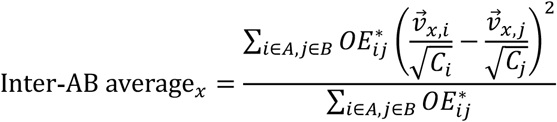

Since *λ*_x_ represents the total weighted sum of squared differences between all interval pairs, we compute the corresponding normalized eigenvalue:

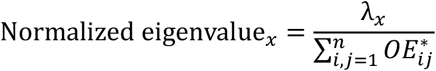

The inter-AB score is then defined as the ratio of these two quantities:

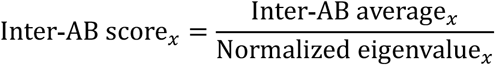

This score quantifies the relative contribution of inter-compartment differences to the total eigenvalue, providing a measure of how well a given eigenvector separates the two compartments.

### Eigenvector Selection with Modified Inter-AB Score

For eigenvector selection, we employed a modified Inter-AB score that incorporates the eigenvalue to balance cut size with compartment separation quality.

We first define the eigenvalue ratio *r*_*x*_ *= λ*_*x*_*/λ*_2_, where *λ*_*x*_ is the x^th^ smallest eigenvalue and *λ*_2_ is the second-smallest eigenvalue. The modified Inter-AB score is then:

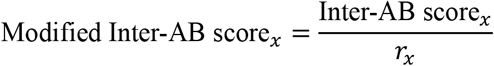

Among the 10 smallest non-trivial eigenvectors (excluding the first trivial eigenvector), the eigenvector with the highest modified Inter-AB score is selected for compartment assignment.

### Orientation Selection

The compartment with higher average intra-interval OE contacts is designated as the A-compartment. We define the average diagonal OE values for each compartment:

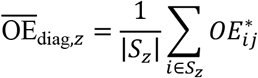

where *z* ∈ {+,-} such that 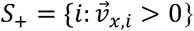 and 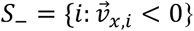 represent the sets of intervals with positive and negative eigenvector values, respectively. The compartment with the higher 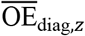 is designated as compartment A, while the other compartment is designated as compartment B.

### Chromosomal Vector Construction

The selected eigenvector 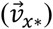 is mapped to a chromosomal vector 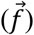 representing the full length of the chromosome. Let *f*(*j*) = *i* denote the mapping from eigenvector index *j* to its respective chromosomal interval *i*:

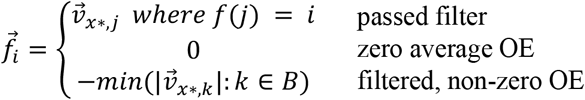

The resulting chromosomal vector is then normalized to unit length.

### Performance Evaluation with Matthews Correlation Coefficient (MCC)

We employed MCC to quantify the agreement between A/B compartment assignments from different datasets. MCC is defined as:

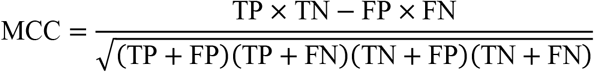

where TP (true positive) represents intervals assigned A-compartment in both datasets, TN (true negative) represents intervals assigned B-compartment in both datasets, FP (false positive) represents intervals assigned A-compartment in the first dataset but B-compartment in the second, and FN (false negative) represents the opposite case. For comparing two A/B compartment assignments, we restricted analysis to intervals that have compartment assignments in both datasets.

We computed MCC values at two levels. For chromosome-level analysis, we calculated MCC between each pair of datasets for individual chromosomes. For genome-wide analysis, we accumulated the TP, FP, TN, and FN counts across all chromosomes before computing the overall MCC value.

### Resolution Selection

The optimal resolution is selected by evaluating whether higher resolution compartment assignments can accurately reproduce lower resolution assignments. For resolution conversion, we determine the ratio between resolutions:

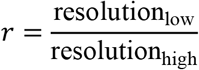

This ratio establishes how many higher resolution intervals combine into each lower resolution interval. This approach requires that one resolution be an integer multiple of the other.

Using the chromosomal vector from the higher resolution 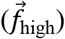, the predicted lower resolution interval value 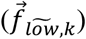 is computed as a weighted average:

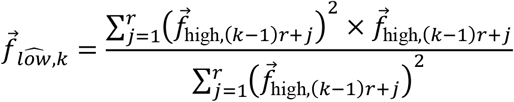

MCC is computed to compare the similarity between the predicted 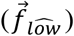 and the assigned compartment 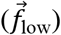.

## Code Availability

The HiC-SCA code is available at https://github.com/iQLS-MMS/hic-sca/.

## References

1. Lieberman-Aiden, E. et al. Comprehensive Mapping of Long-Range Interactions Reveals Folding Principles of the Human Genome. Science 326, 289–293 (2009).

2. Rao, S. S. P. et al. A 3D Map of the Human Genome at Kilobase Resolution Reveals Principles of Chromatin Looping. Cell 159, 1665–1680 (2014).

3. Lafontaine, D. L., Yang, L., Dekker, J. & Gibcus, J. H. Hi-C 3.0: Improved Protocol for Genome-Wide Chromosome Conformation Capture. Current Protocols 1, e198 (2021).

4. Harris, H. L. et al. Chromatin alternates between A and B compartments at kilobase scale for subgenic organization. Nat Commun 14, 3303 (2023).

5. Kruse, K., Hug, C. B. & Vaquerizas, J. M. FAN-C: a feature-rich framework for the analysis and visualisation of chromosome conformation capture data. Genome Biol 21, 303 (2020).

6. Wolff, J. et al. Galaxy HiCExplorer 3: a web server for reproducible Hi-C, capture Hi-C and single-cell Hi-C data analysis, quality control and visualization. Nucleic Acids Research 48, W177–W184 (2020).

7. Open2C et al. Cooltools: Enabling high-resolution Hi-C analysis in Python. PLoS Comput Biol 20, e1012067 (2024).

8. Heinz, S. et al. Simple Combinations of Lineage-Determining Transcription Factors Prime cis-Regulatory Elements Required for Macrophage and B Cell Identities. Molecular Cell 38, 576–589 (2010).

9. Chakraborty, A., Wang, J. G. & Ay, F. dcHiC detects differential compartments across multiple Hi-C datasets. Nat Commun 13, 6827 (2022).

10. Chen, H. et al. Functional organization of the human 4D Nucleome. Proc. Natl. Acad. Sci. U.S.A. 112, 8002–8007 (2015).

11. Ashoor, H. et al. Graph embedding and unsupervised learning predict genomic sub-compartments from HiC chromatin interaction data. Nat Commun 11, 1173 (2020).

12. Kalluchi, A., Harris, H. L., Reznicek, T. E. & Rowley, M. J. Considerations and caveats for analyzing chromatin compartments. Front. Mol. Biosci. 10, 1168562 (2023).

13. Raffo, A. & Paulsen, J. The shape of chromatin: insights from computational recognition of geometric patterns in Hi-C data. Briefings in Bioinformatics 24, bbad302 (2023).

14. Knight, P. A. & Ruiz, D. A fast algorithm for matrix balancing. IMA Journal of Numerical Analysis 33, 1029–1047 (2013).

15. von Luxburg, U. A Tutorial on Spectral Clustering. https://doi.org/10.48550/ARXIV.0711.0189 (2007) doi:10.48550/ARXIV.0711.0189.

16. Luo, Y. et al. New developments on the Encyclopedia of DNA Elements (ENCODE) data portal. Nucleic Acids Research 48, D882–D889 (2020).

17. Durand, N. C. et al. Juicer Provides a One-Click System for Analyzing Loop-Resolution Hi-C Experiments. Cell Systems 3, 95–98 (2016).

18. Durand, N. C. et al. Juicebox Provides a Visualization System for Hi-C Contact Maps with Unlimited Zoom. Cell Systems 3, 99–101 (2016).

19. Jianbo Shi & Malik, J. Normalized cuts and image segmentation. IEEE Trans. Pattern Anal. Machine Intell. 22, 888–905 (2000).

