## Supplementary Materials for "HiC-SCA: A spectral clustering method for reliable A/B compartment assignment from Hi-C data"

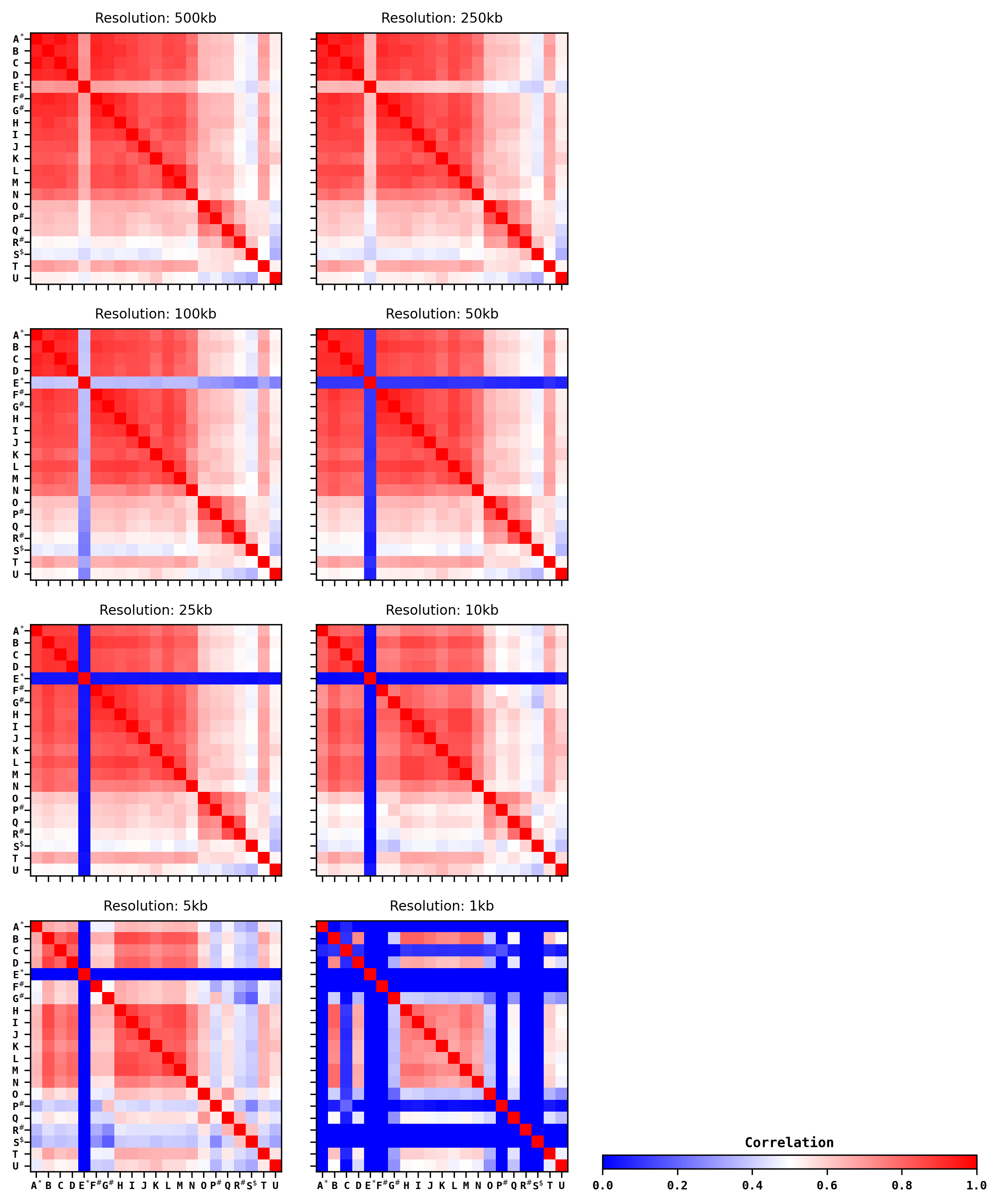

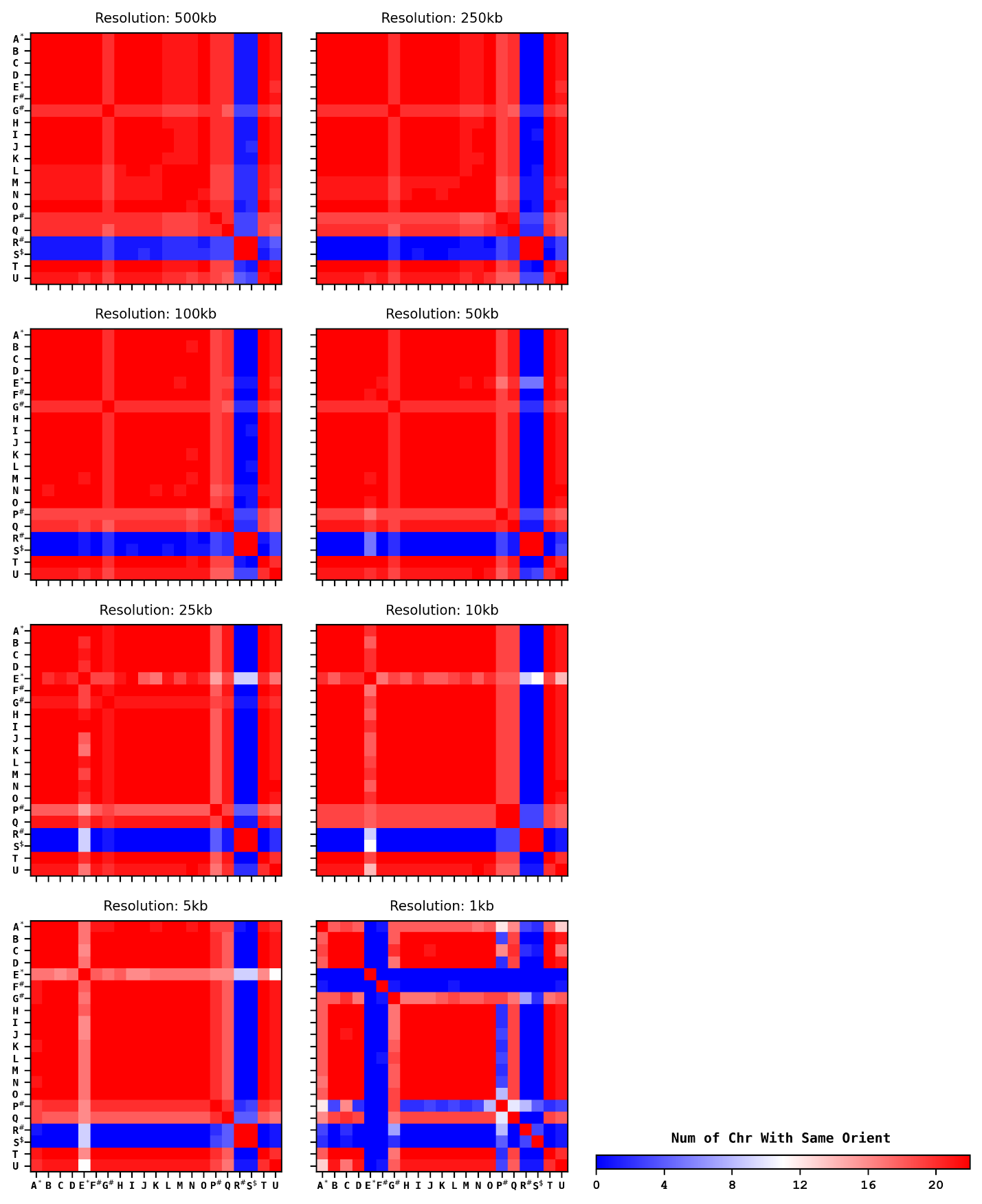

**Supplementary Fig. 1 | Cross-dataset A/B compartment MCC for multiple resolutions.** Cross-dataset MCC matrices for the whole genome at 500 kb, 250 kb, 100 kb, 50 kb, 25 kb, 10 kb, 5 kb, and 1 kb resolution. Dataset markings and the 5 kb resolution data are as shown in **Fig. 2a** of the main text.

**Supplementary Fig. 2 | Cross-dataset A/B compartment orientation agreement.** A/B compartment orientation agreement matrices for the whole genome at 500 kb, 250 kb, 100 kb, 50 kb, 25 kb, 10 kb, 5 kb, and 1 kb resolution. Dataset markings and the 5 kb resolution data are as shown in **Fig. 4d** of the main text.

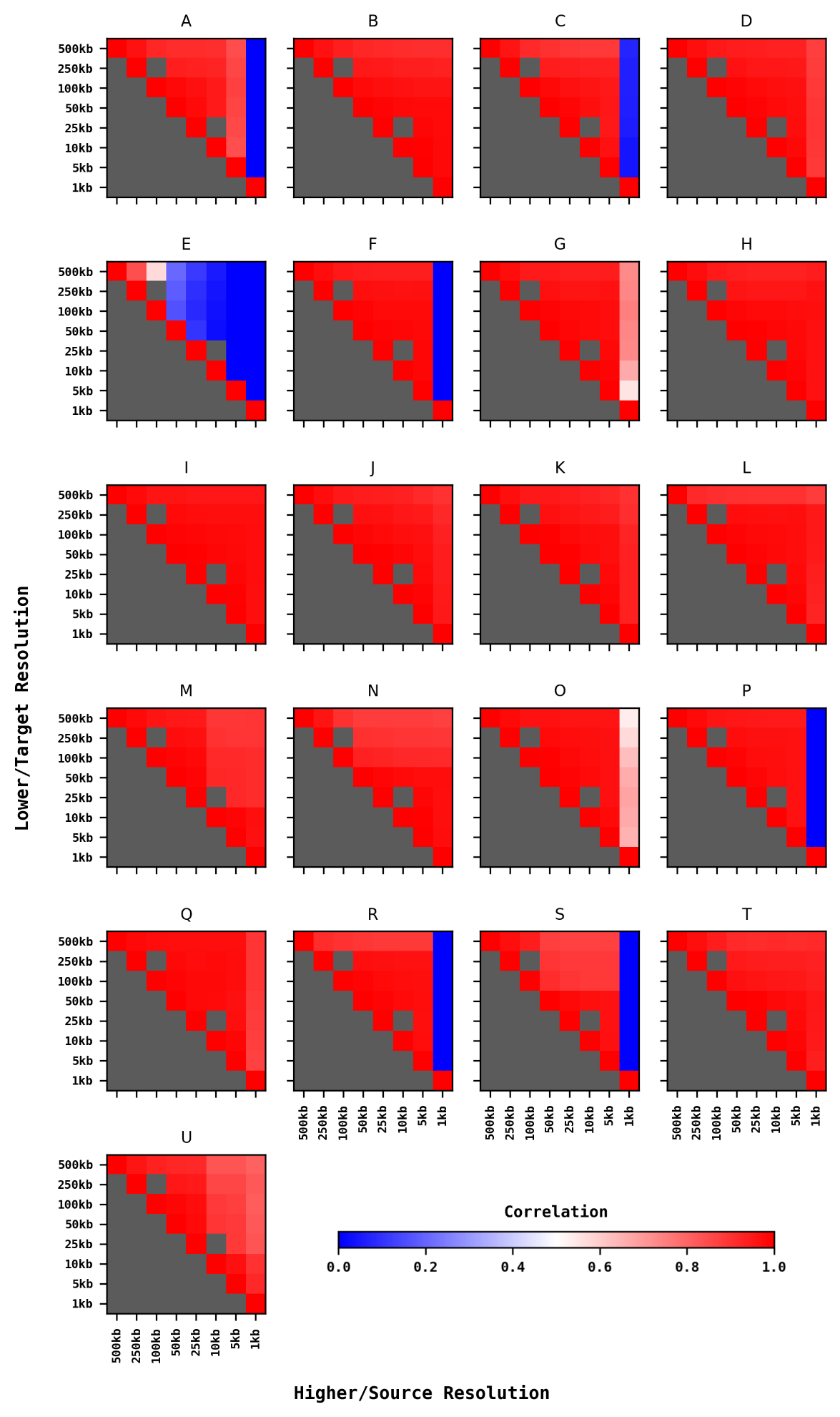

**Supplementary Fig. 3 | Resolution selection evaluation through cross-resolution MCC within the same dataset.** Cross-resolution MCC matrices for all 21 datasets. Dataset G is also shown in **Fig. 6a** of the main text. See **Fig. 6** for details on orientation selection.
